# Automated Feeding and Fermentation Phase Transitions for the Production of Unspecific Peroxygenase by *Pichia pastoris*

**DOI:** 10.1101/2021.07.29.454275

**Authors:** Markus Hobisch, Selin Kara

## Abstract

Fungal peroxygenases are promising biocatalysts for hydroxylation steps in various industry-relevant synthesis pathways. In this application note we describe a bioprocess for the production of unspecific peroxygenase (UPO) in Pichia pastoris. The process was divided in four phases, with different carbon requirements. Precise timing of culture feeding was crucial for optimal cell growth and protein expression. We demonstrate how the automation of culture feeding reduced manual work as well as the risk of process failure due to operator error.

## Introduction

The unspecific peroxygenase (UPO) is a promising tool in synthesis chemistry for enzymatic syntheses based on hydrogen peroxide. Primarily, UPO catalyzes the hydroxylation of aliphatic, cyclic, and aromatic C-H bonds with high enantioselectivity [1,2]. UPO was first discovered in 2004 [3] and its potential for biocatalytic applications was soon apparent, as more and more possible substrates were reported. While there are other enzymes that perform similar functions, they all require elaborate cofactor systems such as P450 monooxygenases, and UPOs only require hydrogen peroxide as a cosubstrate. Besides C-H hydroxylation, the enzyme also catalyzes epoxidation of C-C double bonds and some sulfoxidation reactions. The most widely employed UPO originated from the fungus *Agrocybe aegerita* (*Aae*UPO), was engineered for increased activity and high expression in *Pichia pastoris* and named PaDa-I [4]. The gene is integrated in the yeast’s genome under the control of the AOX1 promoter, which is induced by methanol. Therefore, to grow biomass, glycerol is used as a carbon source, while for protein expression, methanol is required. The process is divided into four phases: Glycerol batch, glycerol feed, methanol batch, and methanol feed phase. Especially during the methanol phase, the process has to be run under carbon limiting conditions to avoid the accumulation of methanol to cytotoxic levels. While it is possible to do this manually, in a fermentation run taking up to seven days, this is laborious and prone to errors that can ruin several days of work. The same is true for the two transitions between batch and feed phases. These components of the process can also be carried out manually, but precise timing and extensive experience are required to avoid stressing the cells without a carbon source or committing to 24/7 monitoring. Therefore developing an automated process was a critical priority.

## Material and Methods

### *Pichia pastoris* strain

The strain *P. pastoris* X-33_pPICZ-B-PaDa-I was kindly provided by Professor Miguel Alcalde, ICP-CSIC, Madrid. The strain carries the UPO gene under control of the AOX1 promotor, which is induced by methanol.

### Culture medium

*Pichia pastoris* pre- and seed cultures were grown in Buffered Glycerol Complex Medium (BMGY) medium. 10 g yeast extract and 20 g peptone are filled up to 800 mL with dH_2_O. After autoclaving 100 mL 1 M Potassium phosphate buffer (pH 6, sterile autoclaved), 100 mL 10% glycerol solution (sterile autoclaved), 2 mL 0.02% Biotin solution (sterile filtrated) are added.

The main fermentation was carried out in basal salt medium, supplemented with PTM1 trace salt solution (**Table 1, 2**).

**Table 1, 2:** Components of basal salt media and PTM1 trace salts solution

| Basal salt media (550 mL) |  | PTM1 trace salts solution |  |
| --- | --- | --- | --- |
| Phosphoric acid (85%) | 24.70 g | Manganese sulfate-H<sub>2</sub>O | 3.00 g |
| Calcium sulfate 2xH<sub>2</sub>O | 0.51 g | Cupric sulfate 5xH<sub>2</sub>O | 6.00 g |
| Potassium sulfate | 10.00 g | Sodium iodide | 0.08 g |
| Magnesium sulfate-7xH<sub>2</sub>O | 8.20 g | Sodium molybdate-2xH<sub>2</sub>O | 0.20 g |
| Potassium hydroxide | 2.27 g | Boric acid | 0.02 g |
| Glycerol | 22.00 g | Cobalt chloride | 0.50 g |
| dH<sub>2</sub>O | 482.00 g | Zinc chloride | 20.00 g |
|  |  | Ferrous sulfate-7xH<sub>2</sub>O | 65.00 g |
|  |  | Biotin | 0.20 g |
|  |  | Sulfuric acid | 5.00 mL |
|  |  | dH<sub>2</sub>O to a final volume of 1 L |  |

### Inoculum preparation

*P. pastoris* was grown on YPD plates containing 100 μg/ml Zeocin for 3 days at 30 °C. A 30 mL preculture (BMGY, 100 μg/ml Zeocin) was inoculated with a colony and grown overnight at 30 °C and 120 rpm in a shaker. A 200 mL seed culture is inoculated with the preculture to an OD_600_ of 0.05 and grown over night as well.

### Bioprocess system and process parameters

Bioprocesses were carried out in a DASGIP^®^ Parallel Bioreactor System with Bioblock equipped with DASGIP vessels (DS1000TPSS, **Figure 1**).Stirring was provided by two Rushton impellers. The bioreactor containing 550 mL basal salt medium was autoclaved and subsequently supplemented with PTM1 trace salt solution (2.4 mL/L). The pH was adjusted to 5.0 with 25 % ammonium hydroxide solution. After DO calibration, process conditions were set to 900 rpm, 30 °C, and 30 L/h air flow. The bioreactor was then inoculated with 50 mL of the seed culture, resulting in a start OD_600_ between 1.0 and 2.5. Antifoam (Antifoam C, Sigma-Aldrich) was added automatically controlled by a level sensor.

**Figure 1:**
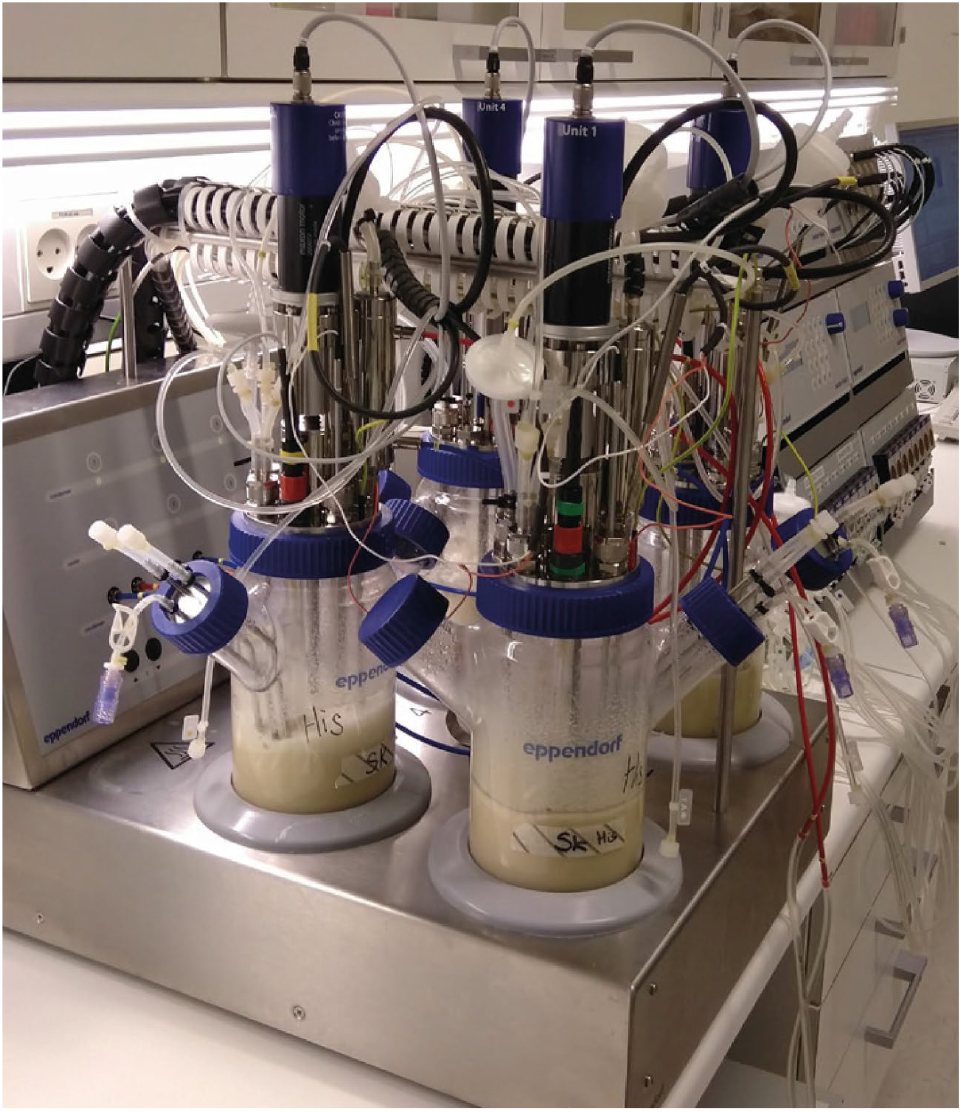
The bioreactor system used in this study

### Bioprocess run

During the whole process the optical density at 600 nm (OD_600_) and cell wet weight (CWW) are monitored. The *Pichia* bioprocess is based on previous publications on UPO expression [4,5] and is divided into four phases:

1. **Glycerol batch phase:** During the glycerol batch phase biomass is generated. During this phase, the dissolved oxygen (DO) will gradually decrease as more cells grow and consume glycerol. When all glycerol and other carbon sources are depleted the DO will spike, triggering the second phase.
2. **Glycerol feeding phase:** During this phase, the culture is fed using pump A of the bioprocess system. The feed solution consists of 25 % glycerol and 2.4 mL/L PTM1 trace salt solution. During this phase, the feed is adjusted automatically to keep the DO at ca. 30 %. This serves to adjust the cells to carbon limited growth conditions and allows formation of more biomass. When a sufficient cell concentration (around 180 g/L) is achieved glycerol feeding is stopped and the third phase, the methanol batch phase, is initiated.
3. **Methanol batch phase:** At the beginning of the methanol batch phase, the temperature is adjusted to 25 °C and methanol (0.5 % of the reactor volume) is slowly added so as to prevent the DO from dropping below 10 %. When all the methanol is metabolized, the DO will spike again and the methanol feed will begin.
4. **Methanol feed phase:** During this phase, the culture is fed musing pump D of the bioprocess system. The feed solution consists of pure methanol and 2.4 mL/L PTM1 trace salt solution. As soon as methanol is added, the enzyme activity in the supernatant is measured with an ABTS assay (see below) until the enzyme activity plateaus, at which time the fermentation is stopped.

### Downstream processing

After cell removal by centrifugation, the supernatant containing the secreted enzyme is concentrated by a factor of 10 using Amicon Stirred Cells with a 10 kDa cut-off membrane. After buffer exchange to potassium phosphate buffer (50 mM, pH 7) the enzyme is frozen by dripping it into liquid nitrogen with the peristaltic pumps from the DASGIP MP8 pump module and stored at −20 or −80 °C.

### Automated feeding based on DO spike

The DO value of the culture medium is influenced by the amount of air supplied to the bioreactor through the bioprocess system’s gassing device and the oxygen consumption of the growing cells. During the bioprocess run, the DO concentration is measured in real-time using a DO sensor. In the course of the bioprocess run it will decrease, reflecting the oxygen consumption of the growing cells. If the carbon sources are depleted, the cells’ metabolic activity and therefore their oxygen consumption suddenly decreases, leading to a spike in the DO concentration. The DO spike therefore indicates substrate depletion and can be used to trigger automated culture feeding. To automate feeding based on the DO spike, a script was implemented in the DASware control 5 software. In the script, the DO spike was defined as the DO sensor rising from a lower trigger point of 25 % to an upper trigger point of 80 %. Upon reaching the upper trigger point, the feed pump was automatically initiated as described in detail in the Results section.

### Analytics

Samples were taken manually to determine OD_600_, cell wet weight, and enzyme activity offline. Determination of cell wet weight 500 μL of cell suspension were centrifuged (7 min, 13,000 rpm) in weighed tubes, and the supernatant was decanted. After another round of centrifugation (1 min, 13,000 rpm) the residual liquid was removed with a 100 μL pipette and cell wet weight determined.

### Determination of Enzyme Activity

The enzyme assay was performed in 100 mM sodium phosphate/citrate buffer pH 4.4, with 0.3 mM 2,2′-azinobis(3-ethylbenzothiazoline-6-sulphonic acid) (ABTS) and 2 mM H_2_O_2_ in semimicro cuvettes (25 °C, at 405 nm, ε=36.8 mM^−1^ cm^−1^).

## Results

After inoculation, the DO started to decrease as expected as more and more biomass was formed, consuming the glycerol (**Figure 2**). It dropped to around 8 %, passing the lower trigger point of 25 %. After around 0.75 days all the glycerol was metabolized and the DO spiked as no more oxygen was being consumed. Upon reaching the upper trigger point of 80 %, the glycerol feed was automatically started with the maximum feed rate defined as 10 mL/h. At first this caused the DO to drop to 10 %, but it soon recovered to 30 % as the feed rate was automatically adjusted to around 8 mL/h. After 40 hours, a sufficient cell wet weight of 188 g/L was reached (**Figure 3**) and the glycerol feed was ended by setting the offline parameter A, which gives the glycerol feed duration from 400 h to 1 h.

**Figure 2:**
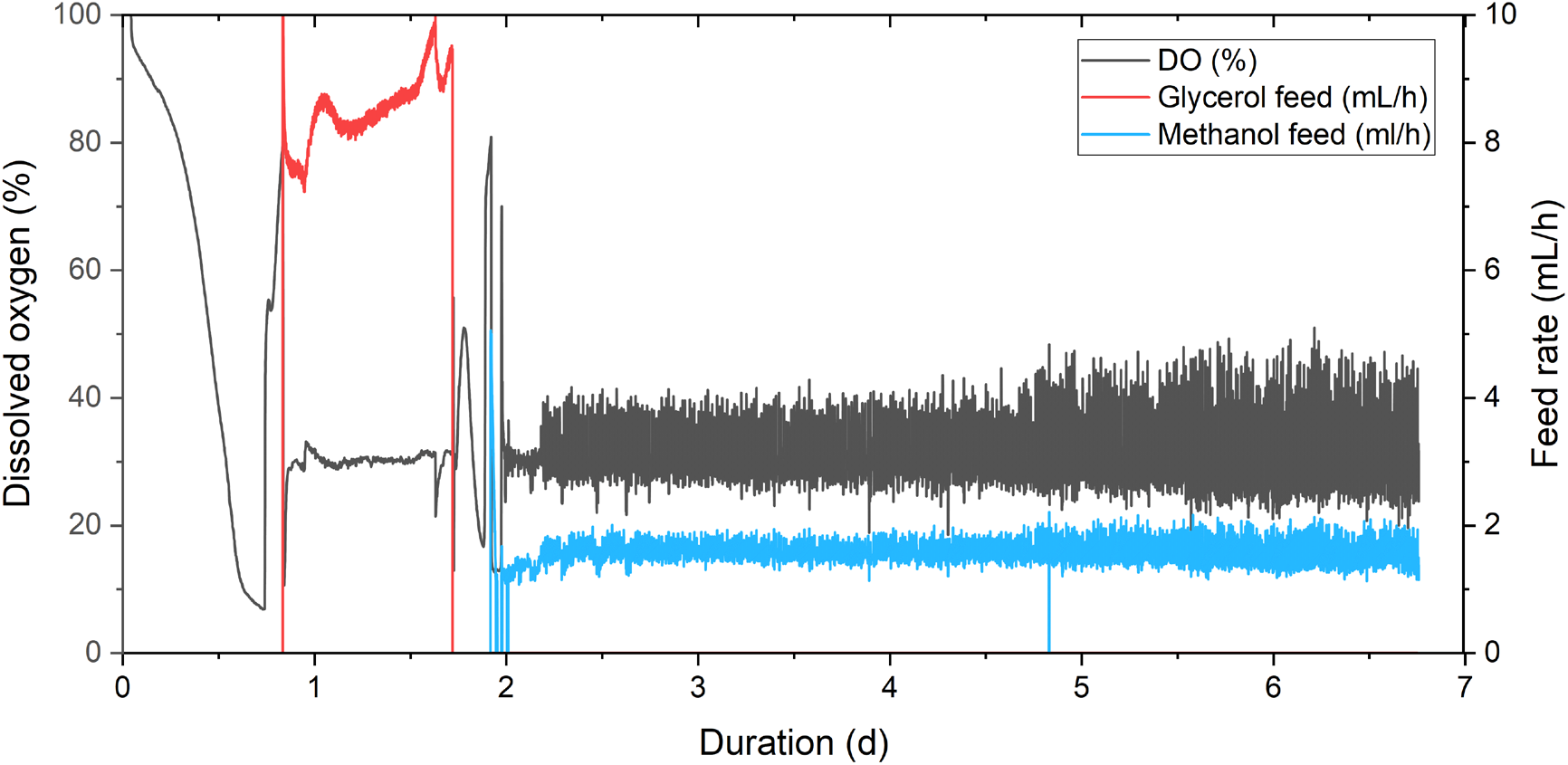
Online parameters of the fermentation comprising dissolved oxygen (DO), glycerol feed and methanol feed.

**Figure 3:**
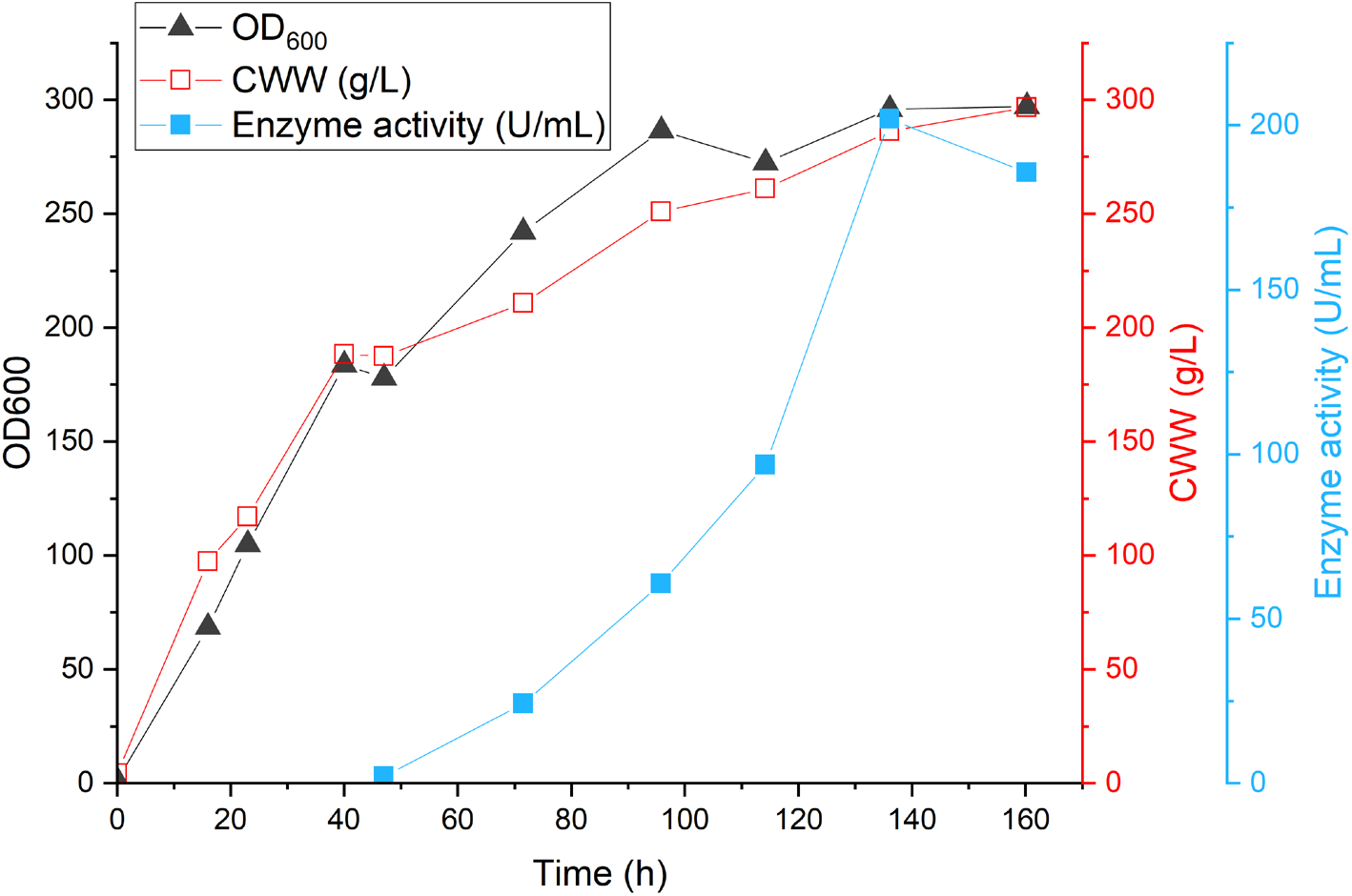
Offline parameters of the fermentation: OD_600_, cell wet weight (CWW) and enzyme activity.

After addition of the methanol for the second batch phase, the DO decreased again to around 15 % (below the lower trigger), but soon spiked as all methanol was consumed, surpassing the upper trigger of 80 % and therefore activating the automated methanol feeding (**Figure 2**). The feed started out with the maximum defined feed rate of 5 mL/h and then settled just below 2 mL/h keeping DO between 20 % and 50 %. Enzyme activity increased to 202 U/mL on the sixth day and slightly decreased to 186 U/mL on the seventh day which was the sign to harvest (**Figure 3**).

## Conclusion

We automated glycerol and methanol feeding in a Pichia pastoris bioprocess for UPO production. To do so, a software script was implemented, which triggered the activity of the feed pumps based on spiking of the DO concentration. It was evident that automated feeding and phase transitions are optimal tools in a fermentation process, especially when dealing with a eukaryotic expression host such as *Pichia pastoris*. The complex process requirements comprising transitions from batch to fed-batch operation and careful methanol feeding over several days are the perfect example for the benefits of automation in a modern bioprocess platform. While a seven day *Pichia pastoris* fermentation is still a demanding and labor intensive process, our proposed modifications represent significant time and labor saving improvements.

## Acknowledgements

This project has received funding from the European Union’s Horizon 2020 research and innovation program under the Marie Skłodowska-Curie grant agreement No 764920.

